# Trace metal availability affects greenhouse gas emissions and microbial functional group abundance in freshwater wetland sediments

**DOI:** 10.1101/515809

**Authors:** Georgios Giannopoulos, Katherine R. Hartop, Bonnie L. Brown, Rima B. Franklin

## Abstract

We investigated the effects of trace metal additions on microbial nitrogen and carbon cycling using freshwater wetland sediment microcosms amended with μM concentrations of copper (Cu), molybdenum (Mo), iron (Fe), and all combinations. In addition to monitoring inorganic nitrogen transformations (NO_3_^−^, NO_2_^−^, N_2_O, NH_4_^+^) and carbon mineralization (CO_2_, CH_4_), we tracked changes in functional gene abundance associated with denitrification (*nirS*, *nirK*, *nosZ*), DNRA (*nrfA*), and methanogenesis (*mcrA*). Greater availability of Cu led to more complete denitrification (i.e., less N_2_O accumulation) and a higher abundance of the *nirK* and *nosZ* genes, which encode for Cu-dependent reductases. We found sparse evidence of DNRA activity and no consistent effect on CO_2_ production. Contrary, net CH_4_ production was stimulated by the trace metal amendments and the Mo additions, in particular, led to increased *mcrA* gene abundance. Taken together, these findings demonstrate that trace metal effects on microbial physiology, which have heretofore only been studied in pure culture, can impact community-level function. We observed direct and indirect effects on both nitrogen and carbon biogeochemistry that culminated in increased production of greenhouse gasses, and the shifts in functional group abundance that we documented suggest these responses may have been mediated through changes in microbial community composition. Overall, this work supports a more holistic consideration of metal effects on environmental microbial communities that recognizes the key role that metal limitation plays in microbial physiology.

## Introduction

Wetland microbes are important for removing many anthropogenic pollutants from surface waters, effectively preventing the contaminants from entering downstream coastal and marine ecosystems. Nitrogen (N) pollutant release (> 60 Tg N y^−1^ globally) to the environment is of particular concern because it is linked to water quality degradation, eutrophication, and increased greenhouse emissions (carbon dioxide (CO_2_), methane (CH_4_), and nitrous oxide (N_2_O)) (Anderson et al., 2010;Tian et al., 2015). In freshwater wetland sediments, the microbial processes of nitrification, denitrification or coupled-denitrification, methanogenesis, and methanotrophy involve N_2_O and CH_4_ as metabolic byproducts, in addition to CO_2_ (Soares et al., 2016). Nitrous oxide is an especially potent greenhouse gas, with a global warming potential (GWP) 298 and 25 times greater per 100 years than CO_2_ and CH_4_, respectively, and with important implications in ozone depletion (Ravishankara et al., 2009;IPCC, 2013). Some microbes can reduce N_2_O to inert N_2_ as part of denitrification, though there is only one enzyme known to facilitate this reaction: the nitrous oxide reductase (NOS)(Thomson et al., 2012). It is estimated that ~17 Tg N y^−1^ is returned to the atmosphere via denitrification in wetlands and terrestrial ecosystems globally, releasing ~48 Tg N y^−1^ to coastal and marine ecosystems (Schlesinger, 2009). Wetlands are equally important for their role in carbon (C) cycling. Despite their relatively small coverage (~10 % of global land area), these ecosystems store large amounts of C and emit considerable amounts of CH_4_ (144 Tg CH_4_ y^−1^)(IPCC, 2014). These net CH_4_ emissions represent a balance between the microbial processes of methanogenesis and methanotrophy.

The microbial production and consumption of N_2_O and CH_4_ is catalyzed by oxy-reductases that utilize metal co-factors (Mo, Cu, Fe) (Glass and Orphan, 2012). For denitrification, the pathway includes the reduction of NO_3_^−^ to NO_2_^−^, catalyzed by a periplasmic or membrane bound molybdenum (Mo) nitrate reductase (NAR) (Schwarz et al., 2009), and the further reduction of NO_2_^−^ to NO by either a non-metal cytochrome c*d*_1_ (Einsle et al., 1999) or a Cu co-factor (Adman et al., 1995) nitrite reductase (NIR), expressed by *nirS* and *nirK*, respectively. NO is further reduced to N_2_O by nitric oxide reductase (NOR), which contains a heme-Fe co-factor (Shiro, 2012). Finally, N_2_O is reduced by a Cu-containing NOS (Rosenzweig, 2000). Similar metal complexes are key in methane cycling. For example, Fe-, Ni-, and Zn-dependent ferrodoxins and dehydrogenases catalyze the early steps of methanogenesis, and all three classes of methanogens utilize a common methyl-coenzyme M reductase (MCR; *mcrA*) to catalyze the terminal steps (Glass and Orphan, 2012). Anaerobic methanotrophs such as members of the NC10 phylum (e.g., *M. oxyfera*) are able to couple CH_4_ oxidation to denitrification (NO_2_^−^-dependent methane oxidation (N-DAMO)) utilizing a Cu-dependent particulate methane monoxygenase (pMMO) and a Fe-rich *cd*_1_ nitrite reductase (Deutzmann et al., 2014;Cheng et al., 2019). Through these sorts of interconnected pathways and processes, the abundance and bioavailability of trace metals in the environment can impact C and N biogeochemistry and help regulate associated greenhouse gas emissions.

Although extensive research has considered the relative contribution of key environmental factors (e.g., pH, NO_3/2_^−^, O_2_, and C/N) in regulating anaerobic microbial processes and specifically denitrification (Wallenstein et al., 2006), our understanding of the impact of metal availability is more limited. Metals in the environment have been traditionally seen as unwanted pollutants, and trace metal accumulation has been linked to toxicity and inhibition of ecosystem processes (Samanidou and Papadoyannis, 1992). For example, concentrations exceeding mg⋅L^−1^ range for Cu, Mo, Fe, Zn and Pb severely inhibited denitrification in soils, sediments, surface water bodies, and waste waters (Labbé et al., 2003;Magalhães et al., 2007;Liu et al., 2016). However, given the dependence of many microbial enzymes on metal co-factors, it is also possible for the opposite problem to occur – wherein function is limited due to an inadequate supply of trace metals. This scenario has received limited study in the environment, but is well documented in case studies with model organisms. In fact, *in vitro* studies have shown that lack of Cu, Mo, or Fe severely inhibits denitrification or methane cycling due to the formation of non-functional enzymes typically lacking the respective metal co-factor. For example, in the soil bacteria *Paracoccus denitrificans* and *Pseudomonas stutzeri*, Cu is required to express to a functional NOS dimer and reduce N_2_O to N_2_ (Granger and Ward, 2003;Felgate et al., 2012;Black et al., 2016). In methanotrophs oxidizing CH_4_ to methanol (CH_3_OH), the switch between a Cu-dependent pMMO or Fe-dependent soluble MMO is regulated by the environmental availability of each metal (Murrell et al., 2000;Bollinger Jr, 2010).

Despite the great progress in understanding metal ecotoxicity and metalloenzyme biochemistry, our knowledge of how trace metal availability affects microbial activity in the environment is generally limited to selected processes (e.g., NH_4_^+^/NO_3_^−^ and CO_2_ assimilation) in aquatic ecosystems (Twining et al., 2007;Glass et al., 2012;Moore et al., 2013;Romero et al., 2013;Schoffman et al., 2016). A broader understanding of how metal availability regulates microbial processes, especially in soils and sediments, is necessary if we are to fully appreciate environmental controls on ecosystem C and N cycling. In this study, we investigated the effects of trace metal additions on the microbial biogeochemistry of freshwater wetland sediments focusing primarily on NO_3_^−^/NO_2_^−^ reduction and greenhouse gas kinetics, and used quantitative (qPCR) to assess changes in the abundance of key functional groups associated with these processes. We hypothesized that greater bioavailability of Mo, Fe, and Cu in the sediments would increase denitrification and that greater Cu availability would reduce N_2_O emissions. We also predicted that C mineralization rates would be impacted by metal additions, specifically higher CO_2_ emissions driven by greater denitrification rates. Furthermore, we expected net CH_4_ emissions to be lower due to a combination of decreased CH_4_ production, since denitrifiers will outcompete methanogens for C substrates once metal limitations are removed, and increased CH_4_ consumption, because of stimulation of methanotrophs through Fe and Cu additions.

## Materials and Methods

### Soil samples

Five soil samples (~350 g each, 5 – 15 cm from surface and ~2 m apart) were collected using a PVC pipe (tube dia. 10 cm and length ~15 cm) from a tidal-fluvial point bar deposit in the Pamunkey River (N 37.557451, W −76.972521) during October 2016. Samples were kept in a cooler on ice during transport to the lab and stored overnight (4°C) until experimental setup (below) and characterization using standard methods (SSSA, 1996). Sediments had 25 (± 4 S.E.) % organic matter (OM), 79 (± 6) % gravimetric moisture content, 0.18 (± 0.05) g.cm^−3^ bulk density, C/N ratio of 11.0 (± 1.5), and pH 6.1 (± 0.3). Porewater, collected from aliquots of soil via centrifugation, contained 0.10 (± 0.03) mΜ NO_3_ ^−^, 0.07 (± 0.02) mM NO_2_ ^−^, 0.16 (± 0.04) mΜ SO_4_ ^2−^, 0.4 (± 0.1) mΜ Cl^−^, 0.3 (± 0.1) mΜ NH_4_ ^+^, 0.11 (± 0.01) mM K^+^, 0.26 (± 0.10) mM Mg^2+^, and 0.29 (± 0.10) mM Ca^2+^.

### Experimental set-up

After large stones and roots were removed, the five soil samples were combined in approximately equal parts to form a composite sample for subsequent experiments. An aliquot (1000 g) of this composite soil was homogenized briefly with an additional 1500 ml of filtered porewater (Cellulose, 10-μm pore-size, Millipore, Burlington, Massachusetts, USA) using a commercial blender (30 s at max speed). Soil slurry aliquots (50 ml) were then transferred to sterile 125-ml glass bottles (Wheaton, Millville, New Jersey, USA). Bottles were crimped with rubber septa and incubated for 3 days in the dark (25°C) to allow residual O_2_ to be consumed by basal respiration. Then, each bottle received a 250-μl aliquot of 2 M KNO_3_ for a final calculated NO_3_ ^−^ concentration of 10 mM. According to the treatment plan (Table 1), replicate (*n*=4) bottles also received aliquots (200 μl) from aqueous stock solutions of ammonium heptamolybdate ((NH_4_)_6_Mo_7_O_24_; 1 mM), iron sulfate (FeSO_4_; 18.5 mM) and copper sulfate (CuSO_4_; 6.5 mM), yielding final concentrations of 28, 74 and 26 μM of Mo, Fe, and Cu respectively. Bottles were vortexed briefly (30 s at 6000 rpm), flushed with N_2_ (60 min), and incubated in the dark (25°C). Gas (5 ml) and soil slurry (1.5 ml) samples were taken anaerobically using a sterile syringe after 0, 6, 12, 24, 48, 72, and 96 h. Following the 96 h sampling, bottles were opened in an O_2_-free chamber and 1 g of wet soil was removed and immediately frozen (−20°C) for future molecular analyses.

**Table 1.**
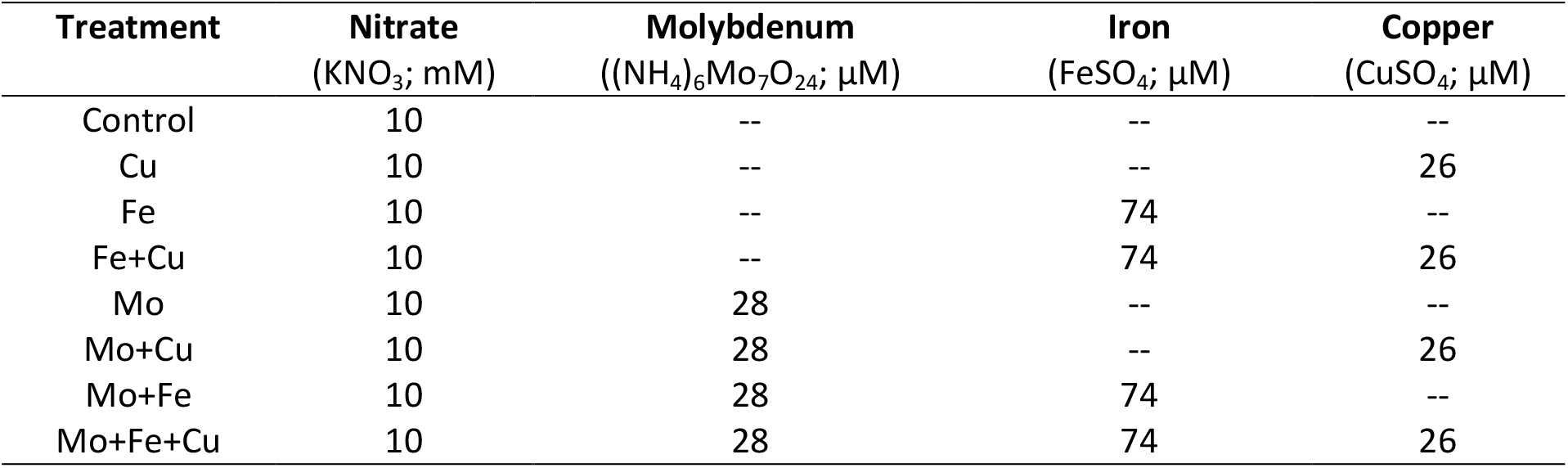
Experimental treatments (*n*= 4).

### Analytical techniques

Gas samples were stored in 3-ml Exetainer vials (Labco, UK) that were previously flushed (N_2_; 5 min) and vacuumed. Concentrations of N_2_O, CO_2_, and CH_4_ were determined with gas chromatography (GC-14A equipped with FID, TCD and ECD; Shimadzu, Columbia, Maryland, USA) as in Morrissey and Franklin (2015). For each gas, total production was determined as the sum of the gas accumulated in the headspace and the gas dissolved in the slurry; the latter was estimated using the relationships described by Heincke and Kaupenjohann (1999). Liquid samples were obtained from each slurry sample by centrifugation (10,000 x *g* for 5 min) and the supernatant was diluted 10-fold, filtered (0.22-μm pore size), and stored frozen (−20°C) until ion analysis. The concentrations of NO_3_^−^, NO_2_^−^, SO_4_^2−^ and NH_4_ ^+^ were determined by ion-chromatography (ICS – 5000+, Dionex, Sunnydale, California, USA) according to the manufacturer’s instructions using columns Dionex IonPac AS17 (2 × 250 mm) and IonPac CS12 (A – 5 µm, 3 × 150 mm) and a Dionex conductivity sensor.

### Molecular techniques

DNA was extracted from frozen soil samples (~1 g wet weight, equivalent to ~0.3 g dry weight) using the DNEasy Kit from Qiagen (Germantown, Maryland, USA) following the manufacturer’s instructions. The retrieved DNA was quantified using a Nanodrop spectrometer (Thermo, Wilmington, Delaware, USA) and visualized on a 1.2 % agarose gel for integrity. The DNA yields were, on average, 42 (± 2) ng⋅μl^−1^. DNA samples were diluted to 3 ng⋅μl^−1^ prior to use in quantitative PCR (qPCR) to determine the abundance of microbial groups typically found in wetland sediments: total eubacteria (*eub-518*), nitrite reducers (denitrification: *nirK* and *nirS*; DNRA: *nrfA*), nitrous oxide reducers (*nosZ*), and methanogens (*mcrA*). SensiFAST™ SYBR® No-ROX Kit Polymerase 2X mix (Bioline, UK) was used according to the manufacturer’s instructions. Primers were purchased from IDT (Integrative DNA Technologies, Skokie, Illinois, USA) and the DNA for the standard curves was extracted from isolates obtained from ATCC (American Type Culture Collection, Manassas, Virginia, USA). Data were collected using a BioRad CFX-384 Real Time System (BioRad, Hercules, California, USA) and analyzed using CFX Manager software (Ver 3.1).

### Estimation of the combined GHG emissions (CO_2_ equivalents)

To compare overall GHG emissions across treatments, we estimated the combined “CO_2_-equivalents” for CO_2_, N_2_O, and CH_4_ based on the 100 y GWP factors (IPCC, 2013). The combined CO_2_-equivalents as g C per bottle were calculated as follows:

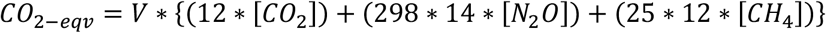

where V is the volume, 298 and 25 is the global warming potential for 100 y per weight basis in CO_2_-equivalents for N_2_O and CH_4_, respectively; 12 the molecular weight of C per mole CO_2_ and CH_4_; 14 the molecular weight of N per mole N_2_O; [CO_2_] and [CH_4_] are the M concentrations for CO_2_ and CH_4_, respectively; and, lastly, [N_2_O] the M concentration of N_2_O per mole N.

### Statistical analyses

Kinetic and molecular datasets were tested for normality (RStudio [*base*]; Shapiro.test). Non-parametric *Kruskal-Wallis* with *Bonferroni* correction statistical tests were applied to assess differences in the mean of ranks between the treatments (RStudio [*agricolae*]). Basic statistics (mean and standard error) were summarized with RStudio [*dyplr*] (Boston, Massachusetts, USA) and plotted with SigmaPlot 14 (Systat Software Inc. San Jose, California, USA). For all statistical tests, *p*≤0.05 was considered significant.

## Results

### The effect of trace metal additions on NO_3_^−^, NO_2_^−^, and NH_4_^+^ kinetics

At the end of the incubation, all treatments had visible gas bubbles on the surface and within the slurry, indicating microbial activity and gas production. We observed yellow and black patches within the slurry possibly due to Fe and residual Mn reduction, indicating anaerobic reducing conditions. Nitrate reduction commenced at the same time for all the treatments (12-24 h, Figure 1), and ~12 mM NO_3_^−^ was consumed by the end of the incubation. At that time, 0.3 mM NO_2_^−^ remained across all treatments while NH_4_^+^ ranged from 0.20 to 0.75 mM (Figure 1). The treatments receiving additional Mo+Fe (*p*≤0.01) and Mo+Fe+Cu (*p*≤0.01) had significantly higher NH_4_^+^ than the control, and an increasing trend in NH_4_^+^ concentration for all treatments was observed towards the end of the incubation.

**Figure 1.**
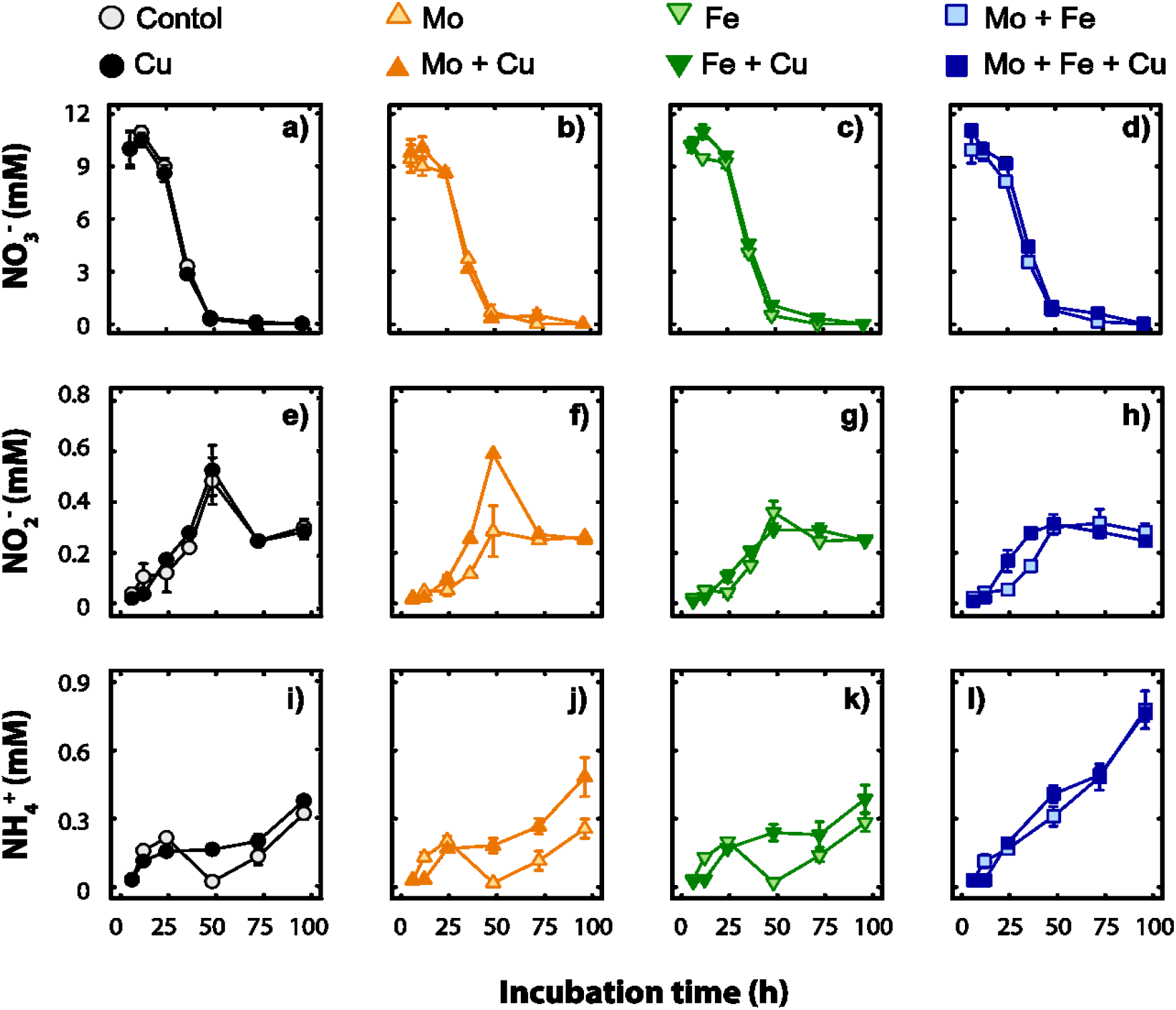
Average NO_3_^−^ (a, b, c, d), NO_2_^−^ (e, f, g, h), and NH_4_^+^ (i, j, k, l) concentrations (mM, n=4, mean ± SE) per treatment throughout the incubations.

### The effect of trace metal additions on N_2_O, CH_4_ and CO_2_ cumulative gas kinetics

Matching the NO_3_^−^ consumption in the slurry, N_2_O emissions increased rapidly between 24 to 48 h (Figure 2.a). Notably, the control, Mo, Fe, and Mo+Fe treatments accumulated significantly more N_2_O than the remaining treatments (Cu, Fe+Cu, Mo+Cu and Mo+Fe+Cu) that all contained Cu. Total N_2_O concentrations reached a plateau between 48 to 96 h, most likely due to exhaustion of the available NO_3_^−^. CH_4_ accumulated quickly in the microcosms (6 to 12 h) and continued to accumulate up to ~6 mM. By the end of the incubation, all metal additions caused an increase in CH_4_ production relative to the control, though the differences were only significant between the control and the Cu, Mo+Cu and Mo+Fe+Cu treatments (Figure 2.b).

**Figure 2.**
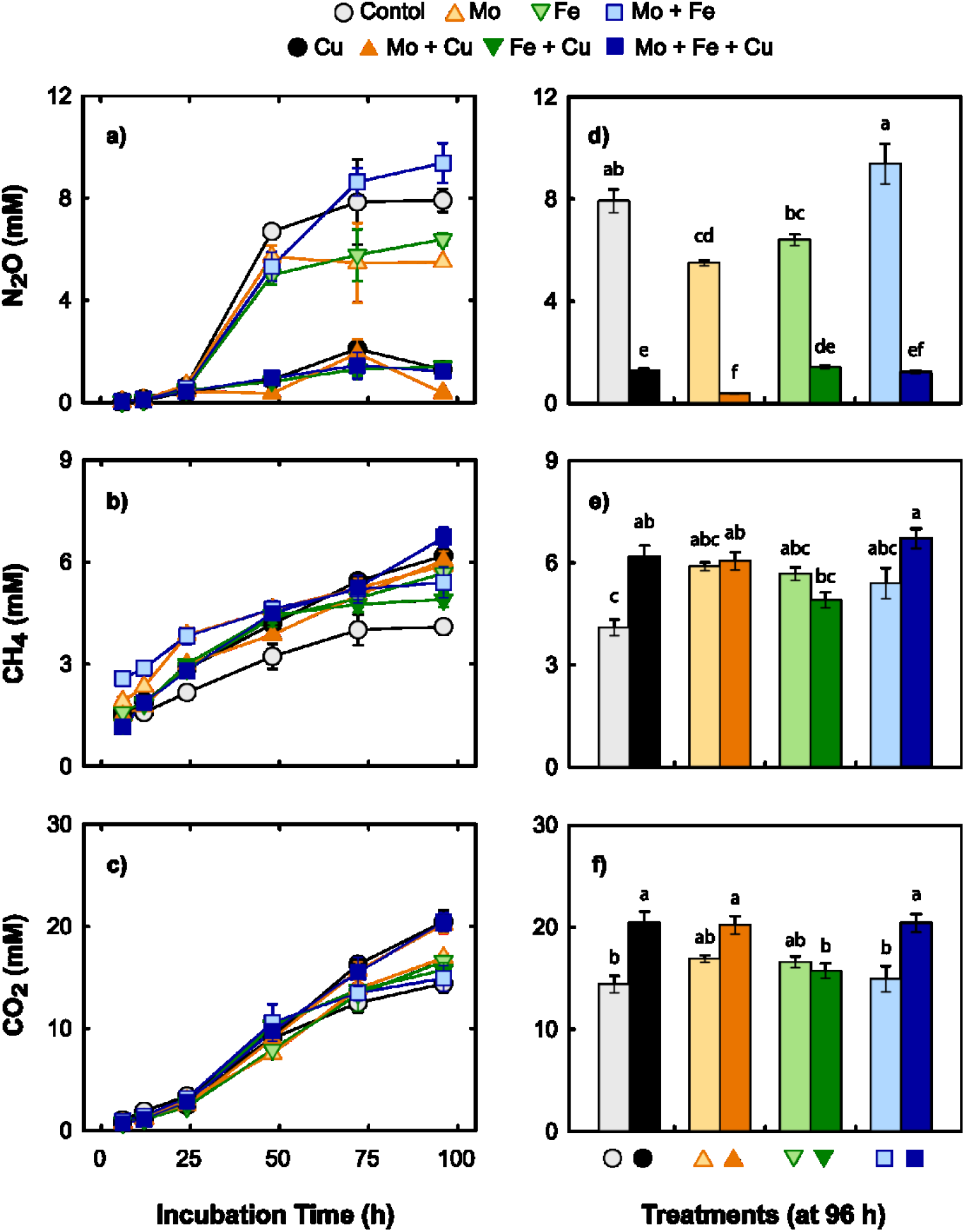
Cumulative gas concentrations of N_2_O (a, d), CH_4_ (b, e) and CO_2_ (c, f), through the incubation and at 96 h, respectively (*n*=4; mean ± SE). Within each bar graph (d, e, f), treatments with the same letter are not significantly different as determined via Kruskal-Wallis and Bonferroni *post hoc* testing.

Cumulative CO_2_ production increased most rapidly between 12 to 24 h, and continued to increase up to ~20 mM CO_2_ at the end of the incubation. There were no significant differences, except between the control and the Cu, Mo+Cu, Mo+Fe+Cu treatments (Figure 2.c).

### The effects of trace metal addition on targeted microbial groups

The average bacterial abundance was 6.7 × 10^8^ *eub* copies⋅g^−1^ sediment dry weight with the Fe, Fe+Cu and Mo+Cu treatments having significantly (*p*=0.005) more *eub* gene copies than the control (*eubacteria;* Figure 3). Trace metal addition affected the abundance of *nirK* (*p*<0.001), *nirS* (*p*<0.001), and *nosZ* (*p*=0.001) denitrifying microbial groups. Reducers of NO_2_^−^ utilizing Cu-NIR (*nirK*) were more abundant in all treatments when compared to the control, though the increase due to Fe addition was not significant. Microbial groups having a cytochrome *cd*_1_-NIR (*nirS*) were relatively more abundant in the Mo and Fe+Cu treatments. The *nosZ* microbial group responsible for the reduction of N_2_O to N_2_ was more abundant in the treatments that received Cu. We observed no effect of trace metals on DNRA NO_2_^−^ reducers (*nrfA*; Figure 3) but found that methanogens were more abundant in the treatments containing additional Mo (Mo, Mo+Cu, Mo+Fe and Mo+Fe+Cu; *mcrA*; Figure 3) when compared to the control.

**Figure 3.**
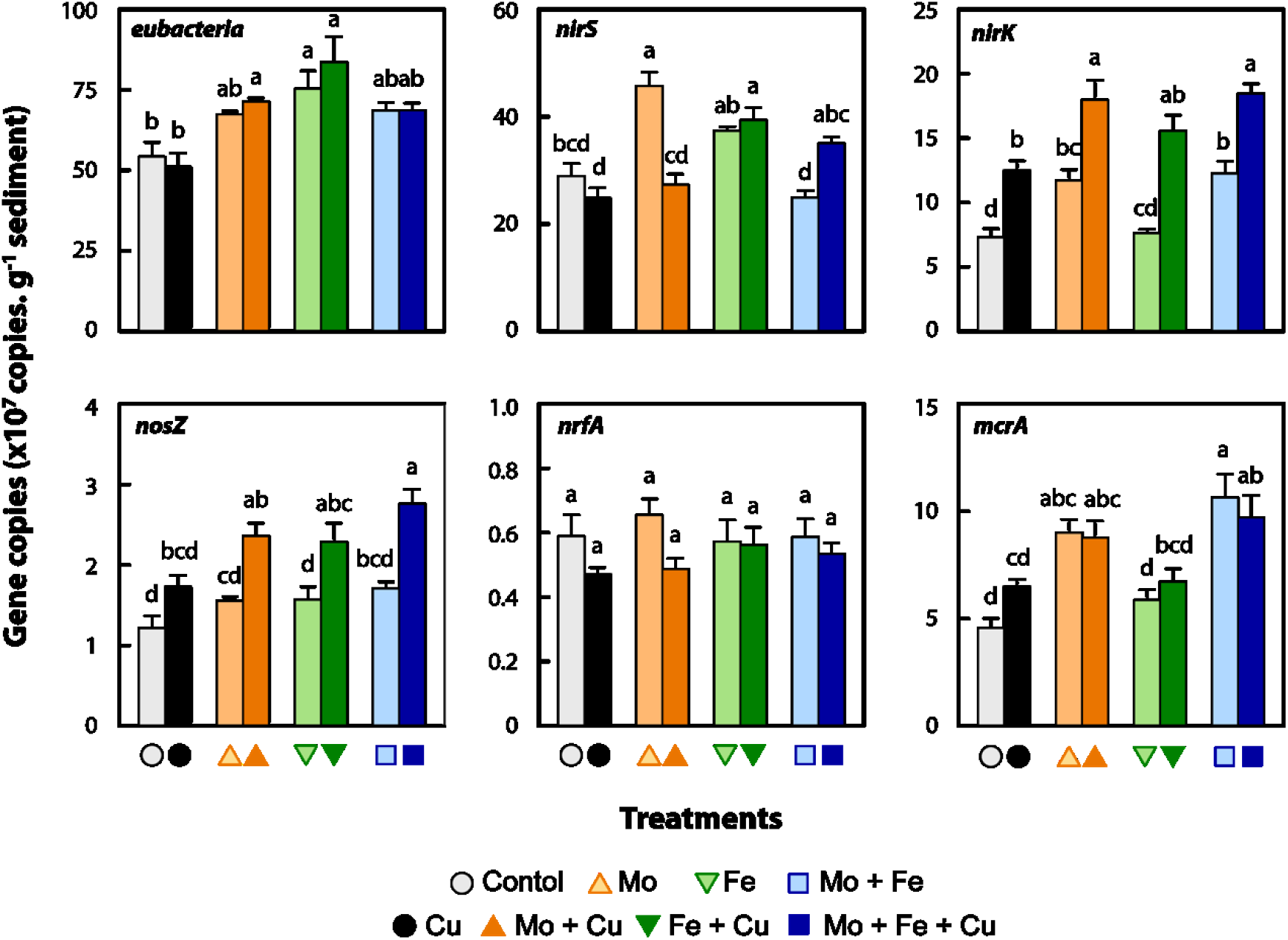
Average gene copies per g sediment (dry weight) at the end of the incubation (96 h; *n*=4; mean ± SE). Within each graph, treatments with the same letter are not significantly different as determined via Kruskal-Wallis and Bonferroni *post hoc* testing.

## Discussion

We observed that trace metal (Mo, Fe, Cu, and their combinations) addition to wetland sediments regulated denitrification kinetics (Figure 1) and greenhouse gas emissions (Figure 2), and altered the abundance of microbial functional groups typically associated with N removal and C cycling (Figure 3). Previous studies have examined the effects of trace metal availability on these processes, often reporting negative effects (Giller et al., 1998;Magalhães et al., 2007;Magalhães et al., 2011;Deng et al., 2018;Keller and Wade, 2018). The range of metal concentrations previously tested far exceeds the levels applied in the current study. The key interpretation of this experiment is that trace metal availability, specifically Cu, enhances the reduction of N_2_O to N_2_, stimulates the production of CH_4_, and has no effect on CO_2_ emissions from wetland sediments. When N_2_O, CH_4_, and CO_2_ were combined and expressed as CO_2_-equivalents, all treatments receiving a Cu addition (Cu, Mo+Cu, Fe+Cu and Mo+Fe+Cu) had significantly lower CO_2_-equivalent emissions (Figure 4), indicating the major role of Cu in anaerobic respiration (denitrification and methanogenesis) as a regulator of nutrient cycling and greenhouse gas emissions.

**Figure 4.**
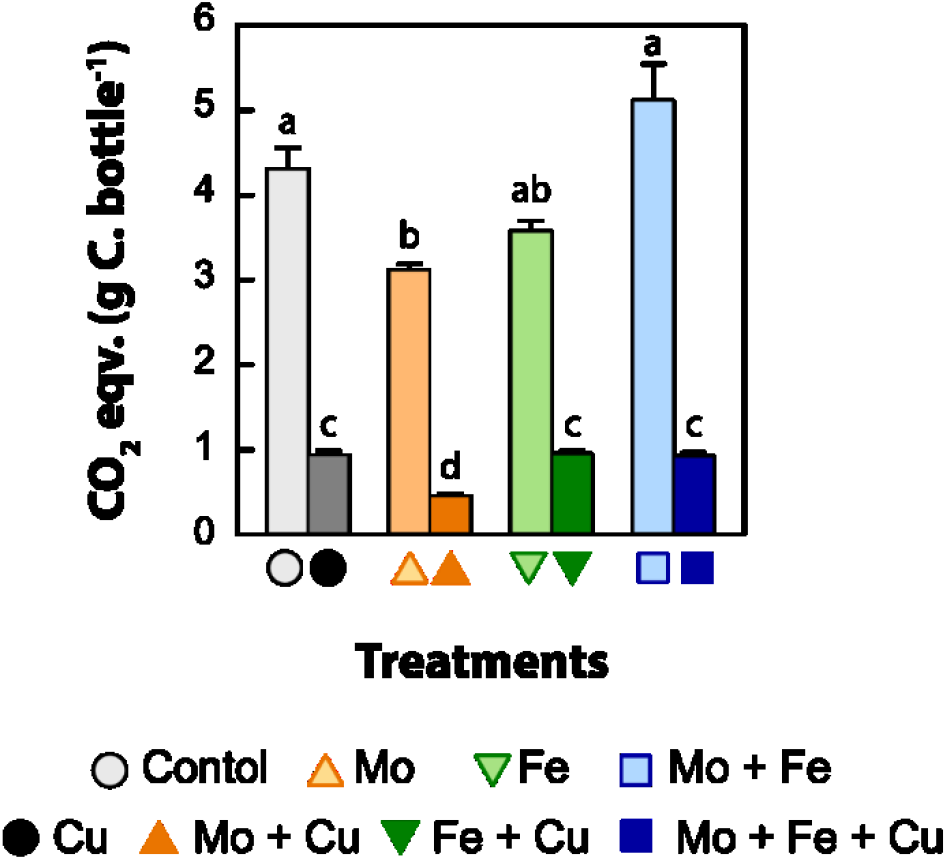
Average cumulative CO_2_-eqv. emissions (100 y^−1^) per treatment at the end of the incubation (96 h; *n*=4; mean ± SE). Treatments with the same letter are not significantly different.

### Denitrification

Denitrification is an anaerobic respiratory process that utilizes NO_3_^−^ as a terminal electron acceptor with sequential reduction: NO_3_^−^ → NO_2_^−^ → NO → N_2_O → N_2_. In our experiments, none of the trace metal additions had an effect on NO_3_^−^ removal (Figure 1), which is somewhat surprising since all bacterial nitrate reductases contain a Mo cofactor at their active sites (Moreno-Vivian et al., 1999). This may indicate that Mo was not limiting to the denitrifiers in this set of soils. Following NO_2_^−^ reduction, the production and consumption of NO is an important denitrification intermediate. Although we didn’t measure NO in our experiments, we assume that it was quickly utilized or reduced to N_2_O due to its potent cytotoxicity at μM levels (Chaudhari et al., 2017;Hartop et al., 2017). In contrast, we saw a dramatic effect of Cu addition. Regardless of what other metals were included, Cu amendments significantly enhanced the reduction of N_2_O to N_2_ (Figure 2) and led to an increased abundance of nitrite (*nirK*) and nitrous oxide (*nosZ*) reductace genes (Figure 3), which both code for Cu-containing enzymes. The especially strong effect of Cu is because the last step of denitrification is catalyzed by a Cu-containing reductase (NOS) and, thus far, no other alternative pathway has been suggested (Thomson et al., 2012). In laboratory experiments, Cu has similarly been found to be a strong driver of denitrifying metabolism. For example, Cu-deficient cells can have more transcripts of *nosZ* and Cu-scavenging genes to compensate for the loss of N_2_O reduction due to depleted Cu availability (Felgate et al., 2012). Also, strains deficient in Cu-transporters and chaperones may be unable to reduce N_2_O to N_2_, even when *nosZ* is expressed and forming a non-functional NOR (Sullivan et al., 2013).

In the treatments accumulating notable amounts of N_2_O (≥ 1 mM), N_2_O cytotoxicity may occur and it can be relieved by the addition of vitamin B12, Co, or methionine (Sullivan et al., 2013). We do not exclude other partial denitrification processes or other anaerobic processes simultaneously co-occurring with denitrification as subjected by the residual N_2_O in the Cu-containing treatments (≤2.3 mM N_2_O). The continuous accumulation of CO_2_ in all treatments firstly, as proxy for microbial activity, suggests against any indirect acute cytotoxicity from NO due to NO_2_^−^ reduction or N_2_O accumulation. Secondly, cumulative CO_2_ emissions indicate that the available electron donor (organic C) was in excess of the added NO_3_^−^. The magnitute of CO_2_ accumulation was predictable; treatments with less residual N_2_O, and thus a more complete denitrifation pathway, had accumulated more CO_2_ by the end of the incubation.

### DNRA

Another possible fate of the added NO_3_^−^ is reduction to NH_4_^+^ via dissimilatory nitrite reduction to ammonium (DNRA). Over the course of the study, we saw small accumulations of NH_4_^+^ (<1 mM; Figure 1), which suggests this process could be happening to some degree, and observed variable effects depending on the type of metal added. The accumulation of NH_4_^+^ was greatest in treatments that contained both Mo and Fe (i.e., Mo+Fe and Mo+Fe+Cu in Figure 1), which may indicate a synergistic response of the enzymes that reduce NO_3_^−^ (all nitrate reductases are Mo dependent) and the DNRA-specific nitrite reductase (*nrfA)*, which includes a multi-heme complex. The addition of either Mo or Fe alone did not result in NH_4_^+^ accumulation. The addition of Cu (added alone, or in combination with Mo or Fe) only led to a modest increase in NH_4_^+^. Overall, the accumulated NH_4_^+^ (<1 mM) was relatively less to the amount of NO_3_^−^ removed (>10 mM) and the abundance of the DNRA marker gene *nrfA* was equally low (Figure 3), so it is unclear whether DNRA was occurring to an appreciable extent in our microcosms. Alternatively, one could postulate that anaerobic organic matter mineralization contributed to the accumulation of NH_4_^+^ and that observed differences due to metal addition were indirect responses. However, if this were the case, one would expect to see a correlation between final NH_4_^+^ concentration and total CO_2_ production, a finding which is not evident. DNRA and N mineralization commonly co-occur in wetland sediments (White and Reddy, 2009), and additional analyses would be necessary to disentangle the two.

### Effect on carbon mineralization

Wetland soils are large C sinks because anaerobic conditions constrain organic matter mineralization and oxidation. The lack of electron acceptors, such as: O_2_, NO_3_^−^, and SO_4_^2−^ in freshwater wetlands, may lead to fermentative and reductive conditions were organic C will be reduced to CH_4_ (Kim et al., 2015;Cheng et al., 2019). In this study, the pattern of CO_2_ emissions primarily reflected the magnitude of denitrification, as discussed earlier. Also, the addition of a more bio-energetically favourable substrate (NO_3_^−^) did not inhibit methanogenesis, nor did the addition of Fe and Cu increase CH_4_ consumption as hypothesized, considering Fe- and Cu-methane monoxygenase pathways; rather, CH_4_ accumulated throughout the incubations. The addition of 10 mM NO_3_^−^, 5 mM NO_2_^−^, and 1 mM N_2_O in anoxic rice field slurries inhibited methanogenesis for 30, 25, and 20 d, respectively, and after that methanogenesis resumed (Klüber and Conrad, 1998). Perhaps, the addition of 10 mM NO_3_^−^ was not sufficient to fully inhibit methanogenesis and/or the ample electron donors from organic matter alleviated any inhibitory effects.

In terms of trace metal amendment, Mo addition enhanced the abundance of *mcrA* genes (Mo, Mo+Cu, Mo+Fe and Mo+Fe+Cu; Figure 3). This was expected because a Mo co-factor is typically required for the formylmethanofuran dehydrogenase (*fwd*) that catalyzes the reduction of CO_2_ to formyl-methanofuran in the first step of methanogenesis by reduction of CO_2_ with electrons from H_2_ (Glass and Orphan, 2012). The subsequent step in methanogenesis is catalyzed by MCR, which is a Ni-containing enzyme. Since we did not add Ni to any in any of our treatments, we assumed that MCR would not change (or would decrease due to compentation with denitrification). Mo addition and the increasing partial pressure of CO_2_ in the bottles during the incubation could have triggered a shift towards a Mo-based methanogenesis pathway. In general, methanogens tend to grow more slowly than denitrifers. It could be that the *mcrA* response was not as strong due to insufficient time to increase in abundance (Enzmann et al., 2018). Another interesting aspect of Cu effects on CH_4_ emissions is the competition between methanotrophs and denitrifiers for the “Cu monopoly” (Chang et al., 2018), which should be further investigated in environmental samples. Thus, the higher abundance of methanogens (*mcrA*) and noticable accumulation of CH_4_ in all metal treatments than the control indicates an important role of trace metal availability in CH_4_ cycling in environmental samples.

### Potential environmental implications

Previous studies have examined the effects of trace metal abundance or addition on C- and N-cycling in sediments, peat, and agricultural soils. Typically, those studies report a negative effect; however, the range of trace metal concentrations in those studies far exceed the levels of the current study. For instance, Keller and Wade (2018) amended peat slurries with ~200 μM trace element solution and saw significant decrease in CH_4_ emission, most likely due to Cu-induced toxicity. Similarly, Magalhães et al. (2007) observed 85% inhibition of denitrification accompanied with accumulation of substantial N_2_O and NO_2_^−^ at 79 μg Cu⋅g^−1^ sediment (equiv. 1200 μM Cu) in estuaries. In a follow-up study, lower denitrification rates due to Cu (~900 μM) were accompanied by a decline in the abundance and *β*-diversity associated with the *nirK*, *nirS*, and *nosZ* microbial groups (Magalhães et al., 2011). A more recent study that tested the effects of Cu pollution (addition of 100 μg⋅L^−1^, ~1.6 μM) on urban freshwater wetland sediments found significantly reduced CH_4_ emissions, but no change in N_2_O emissions (Doroski et al. (2019). This divergent observation could be due to low NO_3_^−^ concentration (18 μΜ) in relation to available C or the presence of substantial amounts of soil Cu (9.4 μg⋅g^−1^ sediment) at the start of the experiment in both control and Cu treatment.

The abundance of bioavailable metals in the environment, typically in their ion forms, is often observed at several magnitute orders less compared to the routinely reported total metal content. Such trivial amounts could be unavailable to microorganisms due to physiochemical interactions and plant uptake (Giller et al., 1998). At neutral and alkaline pH levels, metals tend to be effectively immobilized as inorganic compounds (metal-oxides, -hydroxides and -carbonates). Additionally, organic complexes such humic ligands are known to bind metal cations lessening their availablity to microbes and plants. For example, the particularly low concentrations of dissolved Cu, Fe, and Mo (~5, 1 and 0.2 μg⋅L^−1^, respectively) and the copious dissolved organic C (~80 mg⋅L^−1^) in Siberian peatlands (Raudina et al., 2017) could trigger significant emissions of CH_4_ and N_2_O with climate warming (Basiliko and Yavitt, 2001;Voigt et al., 2017). On the other hand, in temperate salt marshes with reducing conditions, ample SO_4_^2−^ may support abundant communities of sulfate-reducing bacteria that outcompete methanogens and thus suppress CH_4_ emissions, but the H_2_S produced due to SO_4_^2−^ reduction may limit the availability of trace metals through the production of insoluble metal-sulfide complexes with potential implications in N cycling and N_2_O emissions (Gauci et al., 2004;Butterbach-Bahl et al., 2013). In this study, we added 26 μM and 74 μΜ of SO_4_^2−^ as CuSO_4_ and FeSO_4_, respectively, that could potentially have been reduced to H_2_S; however, the initial and final porewater SO_4_^2−^ concentrations remained at comparable levels (average change over time across treatments was only 2.8%, Table 3) and no sulfide odor was detected when microcosms were opened.

**Table 2.** Primers and reaction conditions for qPCR assays

| Target | Primer | PCR Mix (15 µl) | Standard | Reaction Conditions | Reference |
| --- | --- | --- | --- | --- | --- |
| Eubacteria<br>(16S rRNA) | *eub338*<br>*eub518* | 1.2 ng template<br>0.1 µM each primer | *Desulfovibrio desulfuricans*<br>ATCC 27774 | 95°C for 4 min, then 40 cycles of 30 s at 95°C, 30 s at 55.5°C, and 60 s at 72°C. | (Fierer et al., 2005) |
| Denitrifiers<br>(*nirS*) | *cd3aF*<br>*R3cd* | 10 ng DNA template<br>0.1 µM each primer | *Paracoccus dentrificans*<br>ATCC 17741 | 95°C for 4 min, then 50 cycles of 30 s at 95°C, 30 s at 56°C and 60 s at 72°C. | (Throback et al., 2004) |
| Denitrifiers<br>(*nirK*) | *nirKq-F*<br>*nirK1040* | 1.5 ng DNA template<br>0.35 µM each primer | *Pseudomonas nitroreducens*<br>ATCC 13867 | 15 min at 95°C, 9 touchdown cycles of 95°C for 15 s, 68 °C for 60 s, and 81.5°C for 30 s (-1° per cycle for annealing); then 28 cycles of 95°C for 15 s, 60°C for 60 s, and 81.5°C for 30 s. | (Smith et al., 2007) |
| Denitrifiers<br>(*nosZ*) | *nosZ1F*<br>*nosZ2F* | 10 ng DNA template<br>1 µM each primer | *Pseudomonas fluorescens* C7R12 | 15 min at 95°C, 6 cycles of 95°C for 15 s, 67°C for 30 s with a touchdown of -1°C by cycle, 72°C for 30 s, and 80°C for 15 s (acquisition data step); 40 cycles of 95°C for 15 s and 62°C for 15 s, 72°C for 30 s, and 80°C for 15 s; and 1 cycle at 95°C for 15 s and 60°C for 15 s, to 95°C for 15 s. | (Henry et al., 2006) |
| DNRA<br>(*nrfA*) | *nrfA6F*<br>*nrfA6R* | 10 ng DNA template<br>0.3 µM each primer | *Escherichia coli*<br>ATCC 11775 | 50°C for 2 min, 95°C for 8.5 min, and 50 cycles of 20 s at 94°C, 40 s at 54.5°C, and 10 s at 72°C. | (Takeuchi, 2006) |
| Methanogens<br>(*mcrA*) | *mlas*<br>*mcrA-rev* | 2 ng DNA template<br>0.6 µM *mlas*<br>0.7 µM *mcrA-rev* | *Methanococcus voltae*<br>ATCC Strain A3 | 95°C for 5 min, then 50 cycles of 20 s at 95°C, 20 s at 59°C, and 45 s at 72°C. | (Steinberg and Regan, 2009) |

**Table 3.** Initial (0 h) and final (96 h) SO_4_^2−^ concentrations in the various treatments (*n*=4, mean ± SE).

| Treatment | Initial [$\text{SO}_4^{2-}$] $\mu\text{M}$ | Final [$\text{SO}_4^{2-}$] $\mu\text{M}$ |
| --- | --- | --- |
| Control | 4.9 ± 0.8 | 8.5 ± 3.5 |
| Cu | 23.4 ± 2.9 | 20.25 ± 2.0 |
| Fe | 68.9 ± 2.3 | 56.95 ± 8.4 |
| Fe+Cu | 99.2 ± 6.3 | 78.0 ± 33.3 |
| Mo | 15.0 ± 3.8 | 17.5 ± 2.7 |
| Mo+Cu | 21.4 ± 0.9 | 25.4 ± 2.7 |
| Mo+Fe | 68.1 ± 3.9 | 63.5 ± 5.0 |
| Mo+Fe+Cu | 133.0 ± 18.0 | 138.3 ± 21.0 |

Other well-known environmental factors controlling denitrification are O_2_, NH_4_^+^, NO_3_^−^/NO_2_^−^, and C (Thomson et al., 2012). Synthesizing the findings from other chemostat (Baumann et al., 1996;Felgate et al., 2012;Giannopoulos et al., 2017;Hartop et al., 2017;Conthe et al., 2018) and incubation (Koike and Hattori, 1975;Okereke, 1993;Morley et al., 2008) studies, it appears that the relative importance of the aforementioned factors in control of denitrification is O_2_> NH_4_^+^, NO_3/2_^−^> C/N or C quality, with trace metal bioavailability exerting an additional effect on each factor. The present results suggest that the effect Cu exerts on N_2_ and N_2_O emissions from the environment may be more extensive than previously thought. These findings are important not only for understanding fundamental controls on N_2_O production, but also for making predictions about potential future emissions given increasing nutrient and metal loads associated with urban pollution. Interestingly, the effect of trace metal availability on denitrification enzymology is also of considerable interest in the context of agriculture and food production, and it has even been suggested that metal additions could be a possible strategy to mitigate high N_2_O emissions associated with crop production (Richardson et al., 2009).

### Conclusions and future challenges

Our results demonstrate that the importance of trace metal availability in microbial physiology can affect community-level function, including rates of anaerobic respiration, and manifest as differences in ecosystem processes. We found that even short-term (96 hr) exposure to increased concentrations of trace metals led to changes in community structure (functional group abundance) and greenhouse gas emissions (N_2_O, CH_4_, and CO_2_). In particular, our results suggest that trace metal bioavailability may, directly or indirectly, regulate or co-limit denitrification kinetics and favor more active microbial communities that are able to quickly acquire the bioavailable metals and incorporate them in functional oxy-reductases. Disentangling the potential controls of various metals on denitrification, and co-varying effects on C cycling, requires more experiments across a range of metal concentrations and biochemical processes in a variety of different soil types and ecosystems. Future studies should investigate microbial diversity and meta-transcriptomic profiles under comparative treatments to understand and evaluate the ecological importance of trace metal limitation. Such ‘omic studies should be accompanied by high-throughput metabolite or process rate analysis either in the lab or in the field.

## Acknowledgements

We thank Dr. Ioannis Ipsilantis for his comments on an earlier version of this manuscript. We also thank Colin McKenny for assistance in field and lab, Gabriella Balasa for assistance with qPCR, Olivia De Meo for setting up the GC and HPLC procedures, and Dr. Scott Neubauer for equipment access. This work was supported by a grant from the National Science Foundation to Franklin, Brown, and Neubauer (DEB 1355059). This paper is contribution [# to be inserted at time of publication] from the VCU Rice Rivers Center.

